# Extracting herbicide and antibiotic natural products from a plant-microbe interaction system

**DOI:** 10.1101/2023.07.22.550126

**Authors:** Shiyu Pan, Xiaojie Li, Chao Pan, Jixiao Li, Siting Fan, Liner Zhang, Kehan Du, Zhiying Du, Jiayu Zhang, Han Huang, Jie Li, Heqian Zhang, Jiaquan Huang, Zhiwei Qin

**Affiliations:** Center for Biological Science and Technology, Advanced Institute of Natural Sciences, Beijing Normal University at Zhuhai, Zhuhai, Guangdong, 519087, China; Department of Biochemistry and Metabolism, John Innes Centre, Norwich Research Park, Norwich, NR4 7UH, United Kingdom

**Author notes:** **Correspondence** Prof. Zhiwei Qin, Dr. Jiaquan Huang, Dr. Heqian Zhang. These authors have equal contributions (Shiyu Pan, Xiaojie Li).

## Abstract

Plants and their associated microbes live in complicated, changeable, and unpredictable environments. They usually interact with each other in many ways by proceeding in multi-dimensional, multi-scale and multi-level coupling manners, leading to challenges of the co-existence of randomness and determinism, or continuity and discreteness. Gaining a deeper understanding of these diverse interaction mechanisms can facilitate the development of new data mining theories and methods for complex systems, new coupled modelling for the system with different spatiotemporal scales and functional properties, or even universal theory of information and information interactions. In this study, we use a “close-loop” model to present a plant-microbe interaction system and describe the probable functions from the microbial natural products. Specifically, we report a rhizosphere species, *Streptomyces ginsengnesis* G7, which produces polyketide lydicamycins and other active metabolites. Interestingly, these distinct molecules have the potential to function both as antibiotics and herbicides for crop protection. Detailed laboratory experiments combined with comprehensive bioinformatics analysis allow us to rationalise a model for this specific plant-microbe interaction process. Our work reveals the benefits of exploring otherwise neglectable resources for the identification of novel functional molecules and provides a good reference to better understand the system biology in the complex ecosystems.

## Introduction

Soil microorganisms are intricately linked with plants^[1]^. Plant root exudates play a crucial role in attracting beneficial microbes that assist in combatting harmful pathogens and controlling weed growth^[2]^. These interactions primarily involve the exchange of small molecules, including secondary metabolites known as natural products, between plants and microbes^[3]^. While microbial natural products have long served as valuable sources for antibiotics, there is increasing interest in understanding the signaling roles of natural products in inter-species communication^[4]^. Through evolution, plants and microbes have established long-term symbiotic relationships, which can be categorized as mutualism, commensalism, or parasitism, depending on the outcomes such as beneficial interactions, no significant interactions, or defensive responses, respectively^[3]^. Microorganisms that interact with plants are commonly classified as rhizobia, mycorrhiza, endophytes, and epiphytes based on their distribution. It is worth noting that the nature of plant-microbe interactions can vary in different ecological contexts, ranging from beneficial to detrimental and vice versa. In recent years, gram-positive *Streptomyces* species, known for their diverse repertoire of complex molecules, have been observed to interact with plant roots in both the rhizospheric community (the narrow zone of soil surrounding plant roots inhabited by microorganisms) and the endophytic compartment (the intercellular niche within the plant associated with microbes)^[5]^. Numerous studies have demonstrated that plants can selectively recruit *Streptomyces* to their rhizosphere and endosphere, while in response, *Streptomyces* has developed specialized hyphal growth and metabolic networks to facilitate and regulate interactions with plant roots^[6]^.

The biosynthetic gene clusters (BGCs) present in rhizosphere microorganisms encode a diverse array of natural products that serve various functions, including antibiotics, signaling molecules, and potentially undiscovered roles within specific environmental contexts. Our research focuses on identifying and characterizing such molecules through intriguing experimental observations during plant-microbe interactions. By examining these interactions from an evolutionary perspective, we aim to shed light on the long-standing question of why microbes produce secondary metabolites^[7]^. Notably, this perspective can provide insights into why nearly 90% of BGCs remain silent under laboratory conditions, such as stress-free cultivation, and why biosynthetic pathways often yield a range of congeners, precursors, intermediates, and shunt metabolites, each with their distinct pathway specificities, bioactivities, and yields. This understanding contributes to our knowledge of biosynthesis and unveils the underlying mechanisms driving chemical diversity. From a biochemical standpoint, this phenomenon can be attributed to variable reaction rates during biosynthesis. However, the remarkable chemical diversity observed in complex metabolites can be attributed to intricate regulatory networks and pathways that precisely control microbial metabolomic flux, akin to a “closed-loop” system established through a complex yet stable mutualistic relationship. Such unexplored ecological niches hold great potential for discovering new microbes and active natural products, thereby uncovering novel aspects of chemical ecology.

In our recent study, we have examined the capabilities of *Streptomyces ginsengnesis* G7 (G7 hereinafter) isolated from the ginseng rhizosphere. G7 is known to produce a family of type I polyketide alkaloids called lydicamycin (Figure 1)^[8]^. Using this valuable strain, we conducted investigations into two distinct aspects: herbicidal activity and antibiotic properties. In the context of plant-microbe interactions, we made a noteworthy observation regarding the unique behaviour of lydicamycins. Specifically, we discovered that these compounds can function as herbicides by obstructing the auxin transport, a crucial pathway responsible for plant growth regulation. Additionally, our research revealed the presence of uncharacterized metabolites produced by G7, exhibiting potent antifungal activities against crop pathogens such as *Botrytis cinerea*. These findings provide valuable insights into the intricate dynamics of plant-microbe interactions and offer promising avenues for the exploration and development of chemically beneficial compounds or biocontrol agents.

**Figure 1.**
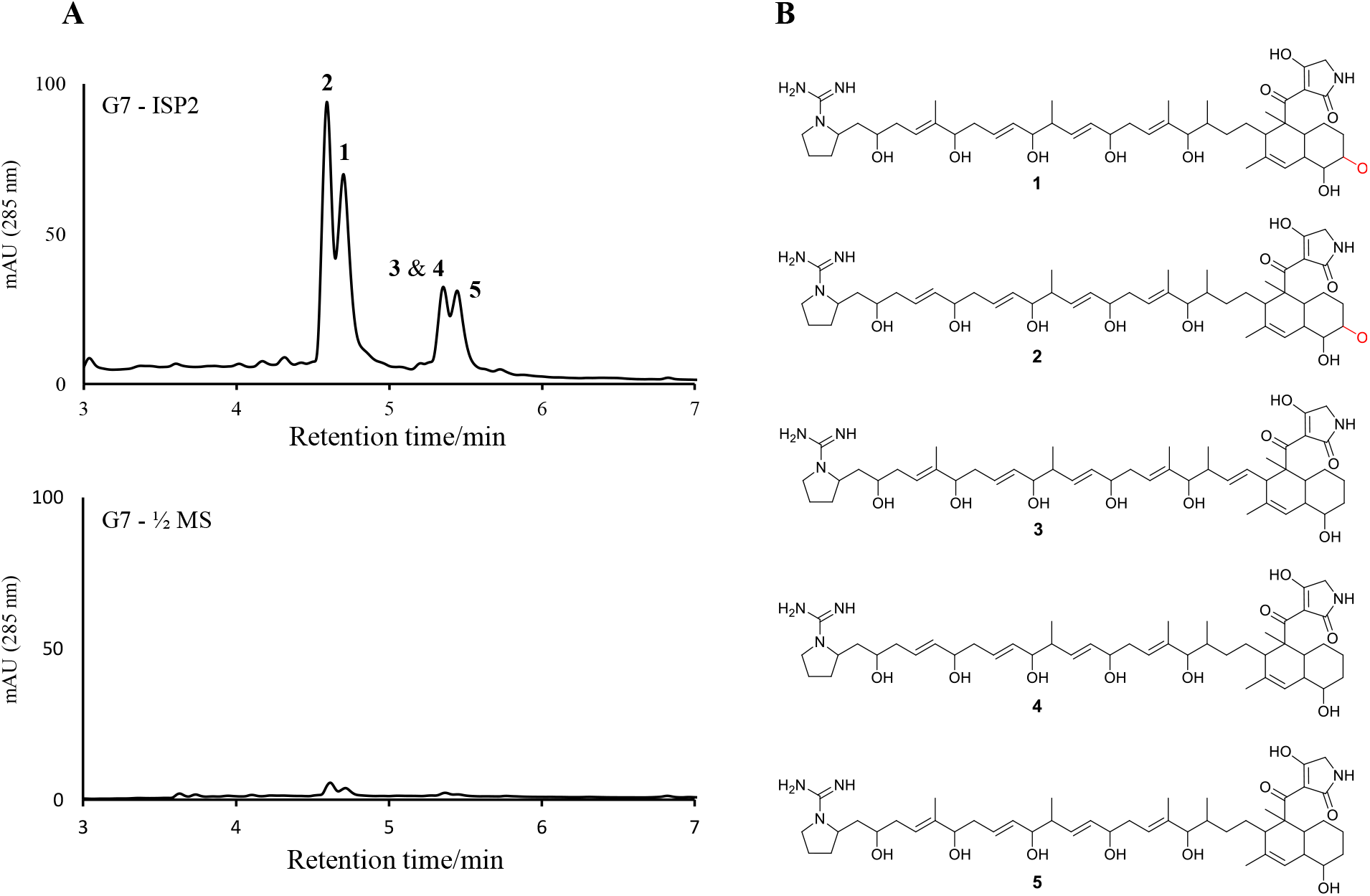
Metabolic analysis. (A) Reconstituted HPLC traces (UV = 285 nm) showing the lydicamycin production from G7 cultivated on different media (ISP2 medium at **top**, and ½ MS medium at bottom). (B) Chemical structures of lydicamycins showing the key difference between **1**-**2** and **3**-**5** in which the hydroxyl groups are highlighted by red.

## Results

### Interactions between G7 and *Arabidopsis thaliana* seedlings reveal inhibitory effects on plant growth attributed to non-volatile lydicamycins

To investigate the potential interaction between G7 and plant seedlings and its impact on plant growth, we conducted extensive experiments using *Arabidopsis thaliana* Col-0 (AT hereinafter) as a model plant. Our goal was to observe visible phenotypic changes induced by G7 and gain molecular-level insights into the underlying physiological and biochemical processes. To achieve this, we developed a protocol in which G7 spores were spread on a standard ½ MS gel plate measuring 10×1 0 cm, covering approximately one-third of the total surface area. After 96 hours of fermentation, AT seedlings were vertically transferred to the aforementioned ½ MS gel plate incubated with the grown G7. To ensure robust statistical analysis, we strategically placed the seedlings in three rows (top, middle, and bottom) with six seedlings in each row (Figure 2A). This approach allowed for the diffusion of any non-volatile substances produced by G7 throughout the entire agar profile, resulting in a concentration gradient that influenced the seedlings on each row through these ‘exogenous materials.’ To account for volatile substances produced by G7, we created a horizontal gap at the top edge of the bacterial inoculation area in the ½ MS agar. This gap prevented the diffusion of non-volatile substances across the agar, while still enabling the free movement of volatile substances within the sealed Petri dish (Figure 2A). All experiments were conducted under identical conditions and repeated at least three times to ensure reliability and reproducibility of the results.

**Figure 2.**
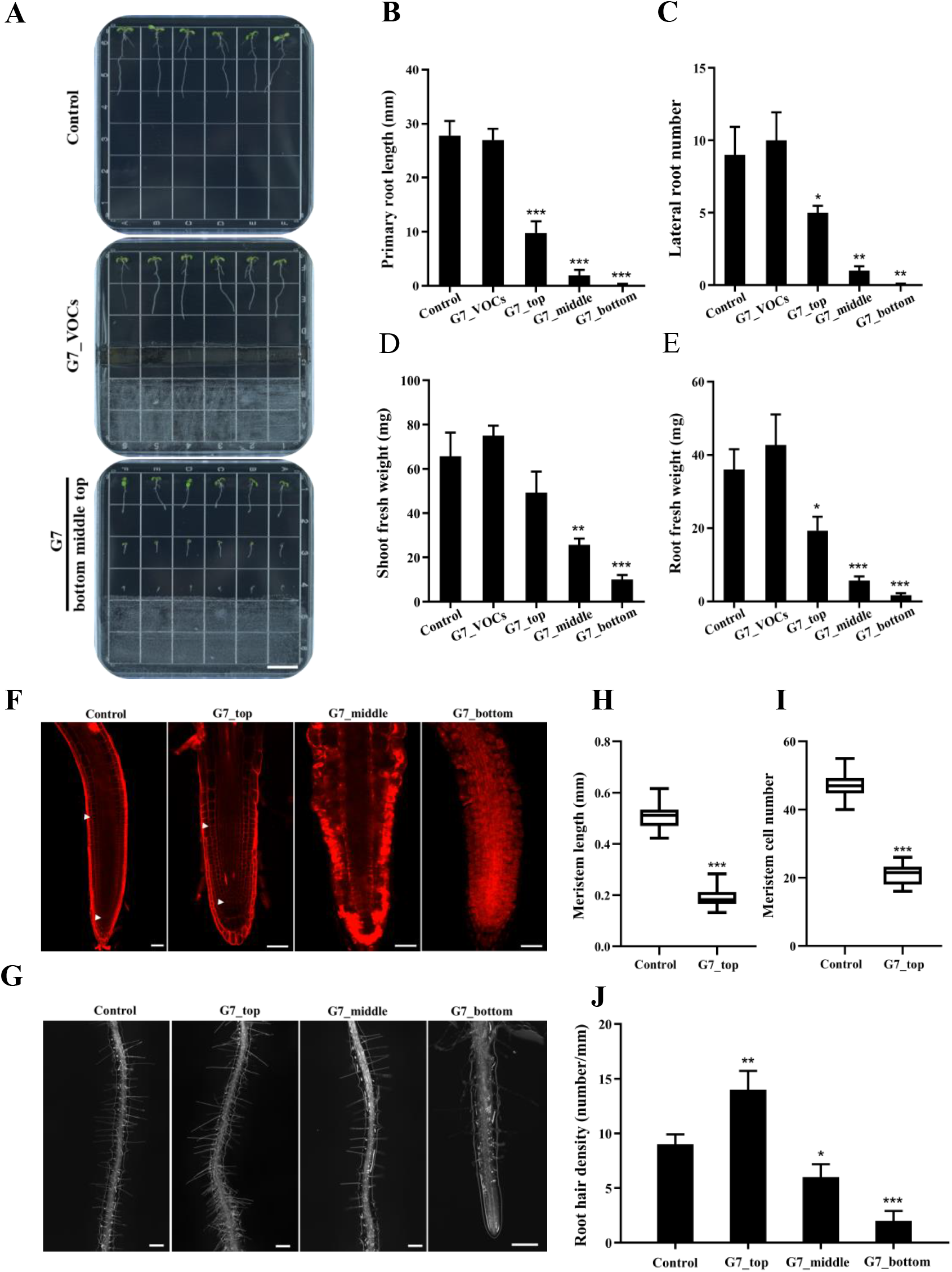
G7 biosynthesizes specific metabolites that exert inhibitory effects on the growth of AT seedlings. (A) Representative images of AT seedlings cultivated on plates without G7 (mock control) and with G7. The positioning of “top, middle and bottom” indicates the varying concentrations of compounds secreted by G7, ranging from low to high. Scale bar: 15 mm. VOCs: volatile organic compounds. (B and C) Quantification of primary root length and lateral root number. (D and E) Quantification of shoot and root fresh weight. Three groups of seedlings were applied, with 15 excised shoots or roots in each group (F) Representative images of primary root meristems of AT seedlings cultivated with and without G7. The white arrows indicate the location of the meristems. Scale bar: 50 μm. (G and H) Quantification of meristem length and cell number. The data represents the median of 30 seedlings. (I) Representative images of root hairs of AT seedlings cultivated with and without G7. Scale bar: 200 μm. (J) Quantification of root hair density. All experiments involving quantification were repeated three times, and consistent results were obtained. All data represents the mean ± SD. SD: standard deviation. T-test: *, *p* < 0.05; **, *p* < 0.01; ***, *p* < 0.001.

The results depicted in Figure 2B-J provided a glimpse of the observed phenotypic changes in terms of primary root length, lateral root number, root hair density, root meristem length and cell number, and fresh weights, suggesting an inhibition of plant growth compared to the mock controls. Intriguingly, these inhibitory effects seemed to be attributed solely to the non-volatile metabolites secreted by G7, as the AT seedlings from the sliced agar plate did not exhibit significant phenotypic alterations (Figure 2A). Since lydicamycins were the only detectable natural products from G7, we speculated that these compounds were responsible for the observed interactions (For more details, see following sections). It is noteworthy that G7 produces lydicamycins on both ISP2 and ½ MS solid agar media, commonly used for microbial and plant growth, respectively, albeit at different titers (Figure 1). Furthermore, we established a calibration curve using purified lydicamycin **1** and calculated the concentration gradient along the ½ MS gel. The results demonstrated a decreasing concentration of lydicamycins from the bottom to the top (data not shown). To further investigate the role of lydicamycin, we repeated the interaction experiment using the mutant strain G7_*Δlyd67*, which we made previously and significantly reduced lydicamycin production by approximately 1000-fold compared to the wild-type strain^[8]^, and found no noticeable phenotypic changes in AT growth in the absence of lydicamycins (Figure 3). Collectively, these findings strongly indicate that lydicamycin plays a role in inhibiting AT growth. However, we cannot rule out the possibility that other undetected metabolites produced by G7 may also contribute to this effect (refer to the antibacterial section). Finally, we evaluated the plant growth inhibition activities of lydicamycins and found that only compounds **1** and **2** exhibited such activities at minimum concentrations of 80 and 100 µM, respectively (Figure 3, Figure S1). This observation is particularly interesting as the genome of G7 shares 96% similarity with *S. lydicus*, a known lydicamycin producer that was previously reported to have no effect on AT plant biomass *in vitro*^[9]^. Thus, the differential behaviour of G7 in our interaction experiments highlights its potential as a biocontrol agent in agricultural applications, and notably, this study represents the first characterization of the plant-interacting activities of lydicamycins **1** and **2**.

**Figure 3.**
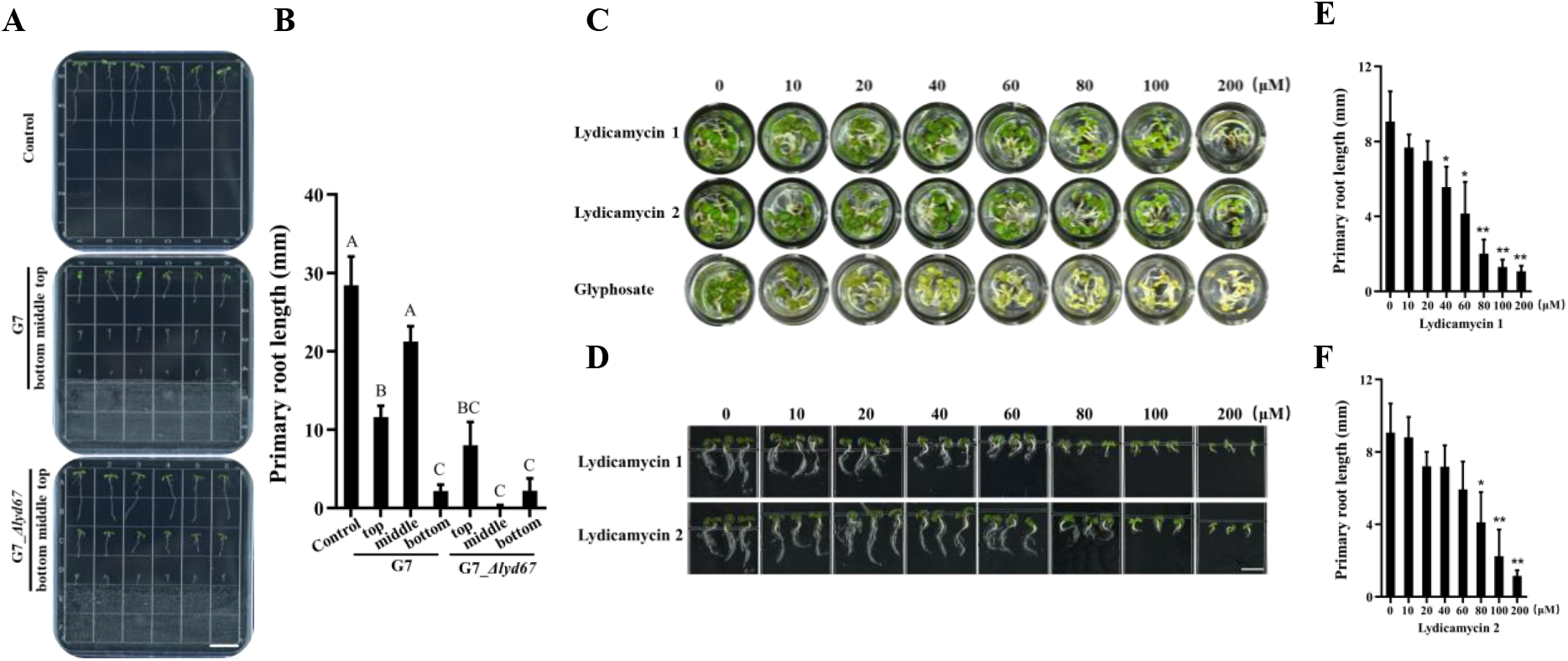
The examination demonstrates that lydicamycin **1** and **2** play an inhibitory role in the growth of AT seedlings. (A) Representative images of AT seedlings cultivated without G7 and with G7 or G7_*Δlyd67*. Scale bar: 15 mm. (B) Quantification of the primary root length in Panel A. (C) The AT seedlings were grown in cells containing various concentrations of **1, 2** and glyphosate. (D) Representative images of AT seedlings treated with **1** and **2** showing in panel C. Scale bar: 5 mm. (E and F) Quantification of the primary root length treating with **1** and **2**. The experiment was repeated three times, yielding similar results.

### Multiomics analysis unravels G7 metabolites primary impact on the root quiescent centre and auxin gradients

At this stage, we conducted a multiomics analysis by profiling the combined transcriptome, proteome, and metabolome, in order to comprehensively investigate the effects of G7 treatment on AT. RNA sequencing (RNA-seq), multiplexed tandem mass tag (TMT) proteomics, and liquid chromatography-mass spectrometry (LC-MS) were employed to analyze the samples treated with and without G7 (Figure 4, Figure S3-4). For the G7 treatment group, we specifically selected the seedlings from the top (referred as TG7) for investigation of differentially expressed genes (DEGs) and differentially expressed proteins (DEPs) at 48 h treatment, as well as differentially expressed metabolites (DEMs) at 96 h treatment. The AT seedlings were collected 48h after being grown on ½ MS gel, on which G7 had already been cultivated for 96 h. This time point was chosen based on our earlier time course study, which indicated that G7 could rapidly produce metabolites typically within 18 h as shown in HPLC (data not shown). We anticipated that at this time point, enough materials from G7 would be present to interact with the AT seedlings and provide insights into the transcriptomic and proteomic profiles. Through preliminary screening, we identified a total of 609 DEGs (|log2FC| > 0.58), 472 DEPs (|log2FC| > 0.26) and 56 DEMs (|log2FC| > 0.58) in the TG7 group. Gene ontology (GO) analysis performed on the DEGs revealed an up-regulated transcriptional pattern associated with root growth, particularly in the maintenance of stem cell populations and root development (Figure 4A). Interestingly, this finding contrasts with the observed restrained root growth in seedlings treated with G7 (see explanations in the next paragraphs).

**Figure 4.**
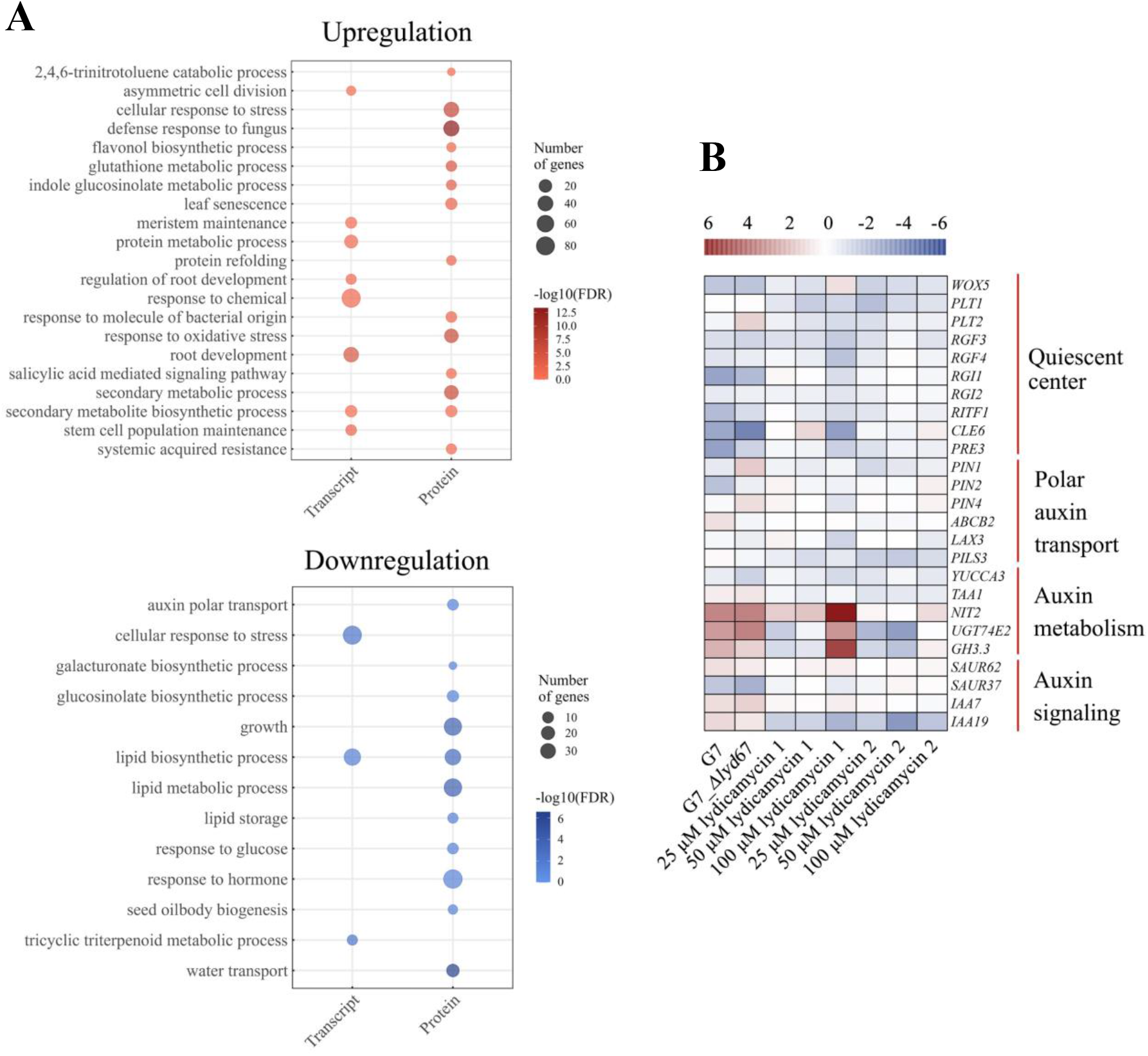
Multiomics profiling studies. (A) Gene ontology (GO) analysis of significantly regulated genes and proteins by G7 compared to the control. Bioinformatics analysis indicates that the G7-induced root phenotype can be attributed directly to the disruption of the distribution of auxin gradient. This is supported by the significant disturbances observed in root development and stem cell population maintenance at the mRNA level, along with the notable downregulation of auxin transport at the protein level. (B) Heatmap illustrating the qRT-PCR analysis of expression levels of QC associated genes, auxin transport, metabolism, and signaling target genes.

Quiescent centre (QC) located within plant roots assumes a crucial role in maintaining stem cell populations and regulating root development^[10]^. Its functions are intricately regulated by modulation of auxin, especially the indole-3-acetic acid (IAA) gradient and distribution^[11]^. Any disturbances in these intricate processes can result in abnormal plant phenotypes and growth patterns^[12]^. In this study, we focused on investigating DEGs and DEPs associated with the TG7 group, specifically examining their connection to the QC homeostasis and IAA metabolism, transport, and signal transduction. Several SAUR genes, including *SAUR9, SAUR37*, and *SAUR62*, were identified among the upregulated DEGs, indicating an increase in IAA levels. Notably, this accumulation raises an intriguing possibility that it may originate from tryptophan-independent auxin biosynthesis, which is supported by the identification of the *NIT2* and *UGT74E2* among the upregulated DEGs and DEPs. However, no significant upregulation of IAA was detected in DEMs, suggesting that free IAA might be conjugated to amino acids since the *GH3*.*3* was upregulated in DEGs. Based on these findings, we speculate that the upregulation of genes expressed in the QC, such as *WOX5, PLT, RGF*, and *RGI*, among the identified upregulated DEGs, indicates the initiation of stem cell maintenance specifically attributed to the auxin maximum within the root tip at the current stage of development. Concurrently, downregulation of certain DEPs related to auxin transport such as PIN2, PIN3 and PIN4 was observed, suggesting that there may be disturbances in the establishment and maintenance of auxin gradients.

To further support our findings based on the transcriptome analysis, we performed RT-qPCR to assess the relative expression of the genes of interest in TG7 seedlings at 96 h (Figure 4B). Initially, we focused on eight QC-associated genes and eleven auxin-associated genes, and isolated mRNA samples corresponding to these genes from the TG7 seedlings. Furthermore, it became evident that at 96 h, the expression of QC-associated genes exhibited downregulation, while the expression levels of IAA transport genes, including *PIN1, PIN2*, and *PIN4*, were gradually downregulated. In contrast to the direct treatment of G7, G7_*Δlyd67* did not result in the downregulation of expression levels for PIN1 and PIN4. This finding suggests that lydicamycin possesses the ability to directly influence the expression of PIN, which is consistent with the observed effects following treatment with lydicamycin **1** at concentrations of 50 and 100 μM. Combined with the above comprehensive multi-omics analysis conducted, it can be inferred that lydicamycins are highly likely to interfere with auxin gradients, then exerted an influence on auxin polar transport, and finally inhibited the root growth.

### Lydicamycin 1 and 2 under certain concentration can stimulate the generation of a lateral root from the meristematic or elongation zone which is previously unprecedented

Inspired by the results so far, we next investigate the molecular mechanisms by which G7 leads to these phenotypic changes in AT (most likely caused by lydicamycins) to reveal the most likely physiological and biochemical processes as reflected by the phenotypes and multiomics during plant-microbe interactions. In order to investigate the molecular mechanisms underlying the inhibition of AT seedling primary root growth by lydicamycin **1** and **2**, we monitored the expression of *CYCB1;1pro:GUS*, which serves as an indicator of G2/M phase in the cell cycle, to detect mitotic activity in the root meristematic tissues and lateral root initiation^[13]^. We analysed the activity of β-glucuronidase (GUS) in 6-day-old *CYCB1;1pro:GUS* seedlings treated with different concentrations of **1** or **2**. Under normal conditions, *CYCB1;1pro:GUS* was expressed in actively dividing cells of the root meristematic tissues. However, after treatment with 25 μM **1** or **2**, GUS staining in the root apex decreased, with **1** exerting a stronger inhibitory effect, whereas at concentrations of 50 μM and 100 μM **1** or **2**, the staining had completely disappeared in the root tip. Furthermore, the most intriguing phenotype observed upon 50 μM **1** or **2** treatment was the emergence of a lateral root in the meristematic or elongation zone (Figure 5A). To the best of our knowledge, such phenotypic changes have not been reported previously.

**Figure 5.**
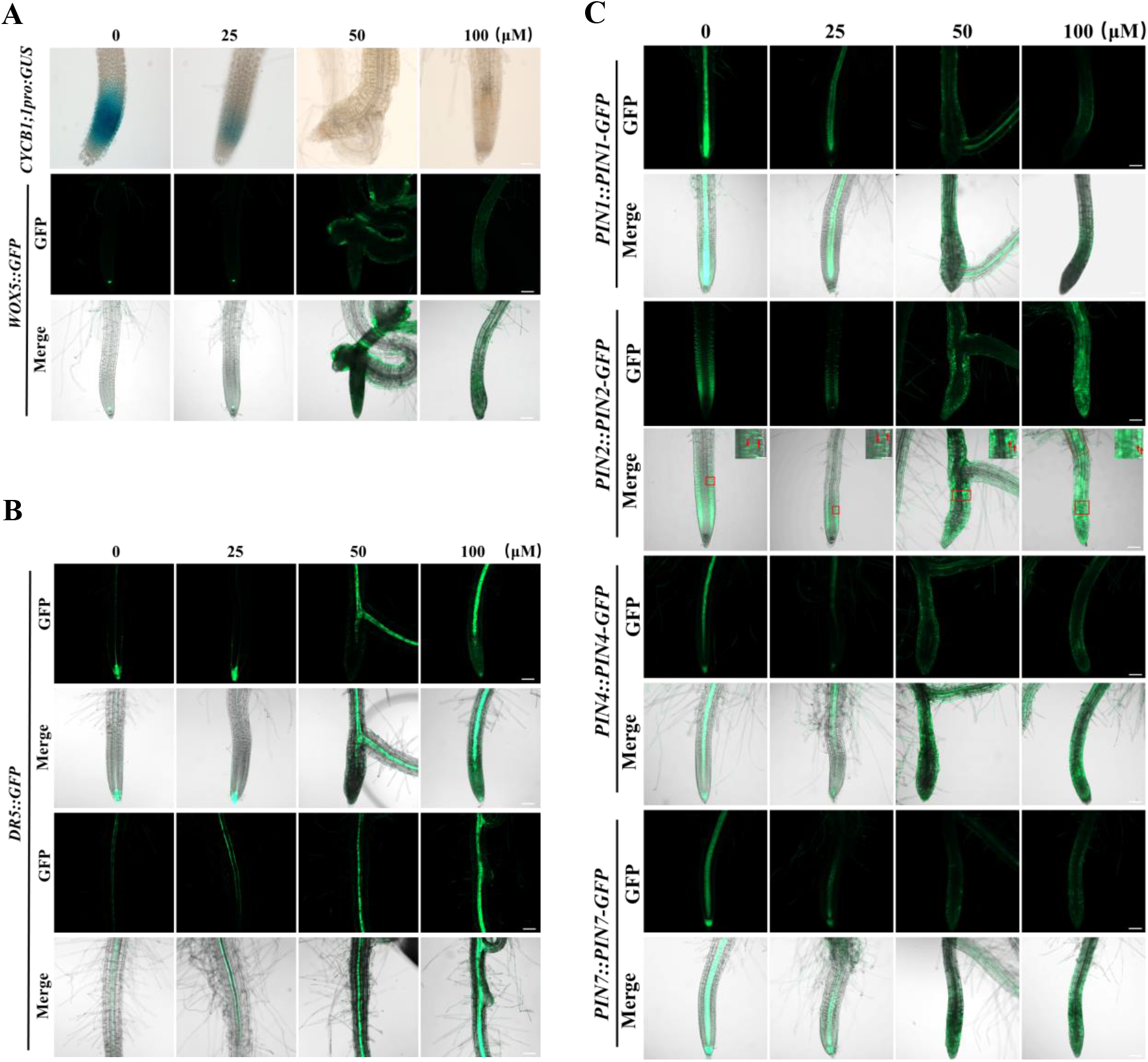
Lydicamycin **1** influences the cell division activity and auxin transport in the root tip of AT seedlings. Representative images of different AT marker lines grown on ½ MS medium supplemented with various concentrations of **1** are shown: *CYCB1;1pro:GUS* and *WOX5::GFP* lines (A), *DR5::GFP* (B), and *PIN1::PIN1-GFP, PIN2::PIN2-GFP, PIN4::PIN4-GFP*, and *PIN7::PIN7-GFP* (C). The scale bar for *CYCB1;1pro:GUS* is 50 μm, while for all others it is 100 μm. Partially enlarged images of *PIN2::PIN2-GFP* are shown with a scale bar of 20 μm.

The discovery of reduced mitotic activity in root meristematic tissues by lydicamycin **1** or **2** prompted us to investigate their potential impact on the stem cell niche, including the quiescent centre and surrounding stem cells^[14]^. To this end, we examined the expression of WOX5 in the root quiescent centre, which is crucial for maintaining the stem cell niche^[15]^. Similar to *CYCB1;1pro:GUS*, the 6-day-old *WOX5::GFP* seedlings exhibited strong GFP fluorescence in the quiescent centre cells of the root meristematic zone. However, after treatment with 25 μM **1** or **2**, GFP fluorescence significantly decreased. Additionally, exposure to 50 and 100 μM of compound **1** or **2** resulted in the disappearance of GFP fluorescence in the quiescent centre cells of the root meristematic zone (Figure 5A).

Based on the observations made thus far, we hypothesize that the intriguing phenomenon could be attributed to the interference of auxin transport. Auxin serves as a key regulator of plant root development and maintains maximum levels around the quiescent centre cells^[16]^. To investigate whether lydicamycins affect root architecture through the auxin pathway, we first utilized AT *DR5::GFP* reporter lines, in which the intensity of GFP fluorescence reflects local auxin abundance and distribution in roots. The 6-day-old *DR5::GFP* seedlings were treated with different concentrations of lydicamycins, and the GFP fluorescence expression was examined. In the absence of treatment (mock), the *DR5::GFP* seedlings exhibited strong GFP fluorescence in the quiescent centre cells, root cap cells, and stele cells of the primary root meristematic zone; whereas in the treatment group, GFP fluorescence in the quiescent centre cells, root cap cells, and stele cells of the primary root meristem vanished (Figure 5B, Figure S2A-B).

The final question raises as which main pathway(s) lydicamycins affect. It has been well understood that it is PIN family that maintain the local gradient of auxin at the root tip which influences root growth by controlling cell positioning, division and expansion^[17]^, we therefore initiated our investigation using *PIN1::PIN1-GFP, PIN2::PIN2-GFP, PIN4::PIN4-GFP* and *PIN7::PIN7-GFP* reporter lines, which were subjected to the same treatment as the aforementioned experiments. The results unveiled that, upon exposure to 50 and 100 μM of **1** or **2**, the GFP fluorescence of *PIN1::PIN1-GFP, PIN4::PIN4-GFP*, and *PIN7::PIN7-GFP* vanished in the primary root tip (Figure 5C). Intriguingly, under normal conditions, *PIN2::PIN2-GFP* was predominantly expressed in the apical epidermal cells and basal cortical cells of the primary root tip. However, upon treatment with 50 μM of lydicamycin **1** or **2**, its expression shifted to the apical epidermal cells and apical cortical cells. Furthermore, at 100 μM concentration, *PIN2::PIN2-GFP* exhibited ectopic expression in the stele tissue and displayed a diffuse distribution throughout the root tip.

It is worth noting that in all reporter lines treated with lydicamycins, the newly generated lateral roots exhibited normal GFP fluorescence. These results indicate that lydicamycin **1** and **2** inhibit the expression of PINs and cause abnormal localization of PIN2. This leads to a decrease in auxin abundance in the root meristematic zone and an accumulation of auxin in the root maturation zone, disrupting the homeostasis of the root stem cell niche. Consequently, the mitotic activity in the root meristematic zone is suppressed, resulting in inhibited primary root growth and enhanced formation of lateral roots at the root tip.

### G7 is a potential biocontrol agent in crop protection

As a final assessment, we aimed to evaluate the potential of G7 as biocontrol agents for crop protection. Initially, we examined the antimicrobial activities of compounds **1**-**5** individually and **6**-**26** as a mixture. Surprisingly, these compounds displayed only mild antibacterial effects (MIC at approximate 20 µM) against reporter strains including *Bacillus subtilis* and *Candida albicans* (Figure 6B), and showed no apparent activities against the crop pathogens *Botrytis cinerea, Fusarium graminearum, Ustilaginoidea virens*, and *Magnaporthe oryzae*. However, in agar-plug bioassay tests, G7 exhibited potent inhibition of the aforementioned pathogens (Figure 6B), indicating that G7 must produce other unidentified metabolites responsible for this activity. To further validate our hypothesis, we employed the mutant strain G7_*Δlyd67* to repeat the above antimicrobial experiments. The results showed that this mutant strain did not display activity against *B. subtilis* and *C. albicans*, but it still exhibited inhibitory effects on the plant pathogens (Figure 6B). Although our attempts to isolate the antifungal compounds were unsuccessful due to undetectable UV and ionic signals, we remain committed to resolving this issue in future investigations. It is worth mentioning that a recent study demonstrated that rhizosphere bacteria employ indole-3-acetic acid (IAA) as an inter-species signaling molecule to enhance the production of active secondary metabolites, including antifungals^[9b]^. In light of this, we tested the antifungal activities of G7 in the presence of IAA (0.1 mg/mL), but found no significant enhancements were observed (data not shown), indicating a distinct regulatory role for antifungal biosynthesis in G7.

**Figure 6.**
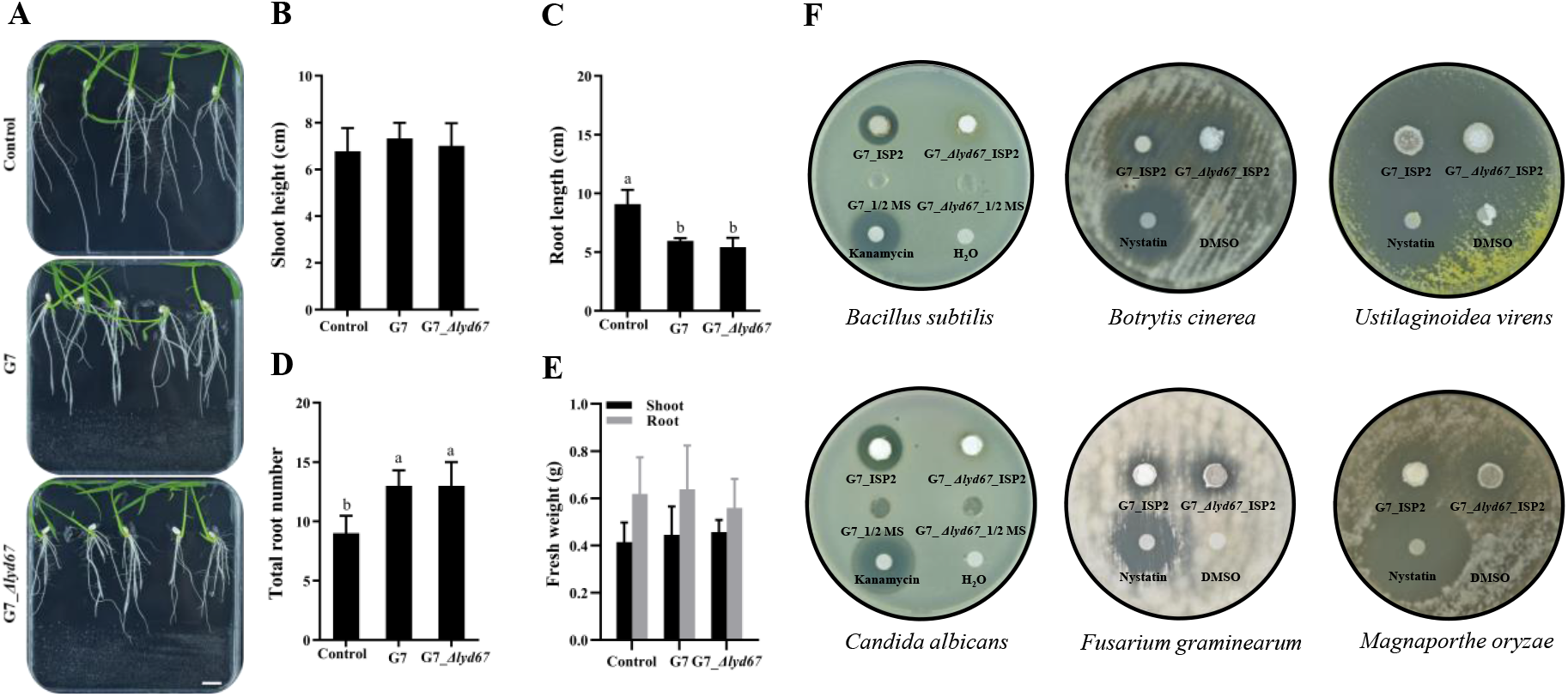
Evaluation of G7 as a biocontrol agent. (A) Representative images of rice seedlings grown without G7 and with G7 or G7_*Δlyd67*. Scale bar: 13 mm. (B to D) Quantification of shoot height, root length, and total root number. (E) Quantification of fresh weight of shoot and fresh roots showing in panel A. Three groups of seedlings were used, with 10 excised shoot and roots in each group. (F) Agar-plug bioassay tests of G7 and G7_*Δlyd67* conducted using a series of laboratory reporter strains and crop pathogens. Kanamycin and nystatin were used as antibacterial and antifungal controls, while water and DMSO as negative controls, respectively. The experiments mentioned above were repeated three times. All data represents the mean ± SD. T-test, *, p < 0.05.

Next, we conducted similar plant-microbe interaction experiments using monocot rice to assess the effects of G7 and G7_*Δlyd67* on crop seedlings. In this study, we focused on evaluating shoot and root parameters as indicators of the interaction. Root systems play a crucial role in nutrient absorption, amino acid biosynthesis, and hormone regulation, which are closely associated with plant growth, development, and yield. We inoculated G7 and G7_*Δlyd67* on ½ MS gel for 96 hours of fermentation, followed by the transfer of 2-day-old crop seedlings onto these plates for an additional 13 days of growth. Comparing with the control groups, we observed that both G7 and G7_*Δlyd67* treatments led to a reduction in root length (approximately 37%) but an increase in root number (approximately 31%), and there was no significant difference between the effects of G7 and G7_*Δlyd67* in this regard (Figure 6A). Interestingly, no significant differences were observed in the shoots and roots of rice seedlings between the treatment groups. Notably, root length, biomass, and root number are critical parameters that reflect water and nutrient absorption by rice comprehensively. Using the rice variety Nipponbare as a reference, we found that G7 metabolites increased the number of roots per rice seedling while decreasing root length. However, it had no effect on the overall growth, including the fresh weight of aboveground and underground parts, as well as the shoot height of rice seedlings. These results indicate that although G7 metabolites inhibited root elongation in rice seedlings, they did not have a significant impact on overall growth. Furthermore, we observed no significant differences between the effects of G7 and G7_*Δlyd67* on rice seedlings, suggesting that lydicamycins did not influence the growth of rice seedlings.

It is worth noting that, in a recent study, it was demonstrated that *Streptomyces* sp. NEAU-S7GS2, a species isolated from the root of *Glycine max* and its rhizosphere, possesses antifungal properties and promotes plant growth. Through genome mining, specific genes associated with phosphate solubilization, auxin biosynthesis, and the production of 1-aminocyclopropane-1-carboxylate (ACC) deaminase, glucanase, and α-amylase were identified^[18]^. Notably, the plant hormone indole-3-acetic acid (IAA) was detected in NEAU-S7GS2 cultures when cultivated in the presence of _L_-tryptophan. In contrast, G7, the focus of our study, did not exhibit robust growth in the presence of tryptophan, and no significant levels of IAA were detected in its extracts under our laboratory conditions (data not shown). These findings suggest that G7 represents a distinct species and differs from NEAU-S7GS2, despite their high similarities.

## Discussion

Plant-microbe interactions are intricate and dynamic bioprocesses, and understanding how nature stimulates microbial responses to produce specific natural products has always been a fascinating topic. Despite numerous strategies devised to activate these important biosynthetic gene clusters (BGCs), there is no universal approach that works optimally for all cases^[19]^. Furthermore, the complexity is compounded by the fact that microorganisms typically inhabit highly diverse and intricate microenvironments or niches, making it challenging to replicate the multiple synchronized conditions found in nature within a laboratory setting^[20]^. In this study, our focus was on G7, a rhizosphere-dwelling species of significant interest isolated from ginseng plant rhizosphere. We demonstrated its unique protective role through the production of lydicamycins and other unidentified natural products with herbicidal and antibiotic properties. Notably, among all fermentation conditions, lydicamycin **1** and **2** were consistently observed as the most abundant metabolites detected by mass spectrometry or chromatography. This suggests that certain congeners may be absent in specific scenarios, such as different culture media or cultivation times. This observation is a common pattern encountered in the process of natural product discovery and is likely influenced by selective pressures^[20a]^. It is currently unclear why G7 does not (or under limit of detection) produce lydicamycins other than **1** and **2** on ½ MS medium the way it does in ISP2 medium. One plausible explanation is that ISP2 medium, being generally richer in nutrients for microbial growth, may modulate the intricate regulatory network of lydicamycin biosynthesis.

This is also reminiscent of another interesting consideration that plant species can recruit beneficial microorganisms for protection, and in response, BGCs in these microorganisms evolve towards beneficial symbiosis. As a result, during long term of evolution, selection has chosen the microorganisms that are capable of producing highly effective and potent chemicals at low cost. This finding is consistent with our observations that G7 produces diverse lydicamycins during a quick fermentation time and repeated metabolic analysis of the crude extracts from bacterial fermentation. Furthermore, as observed, only two (**1-2**) lydicamycin congeners have interactions with AT at certain concentrations (> 40 μM), while the remaining (**3-5**) appear to be neither antibacterial/antifungal, nor herbicidal, and thus we speculated that G7 has evolved its lydicamycin BGC to primarily facilitate the production of congeners **1**-**2**. From this perspective, the chemical structures of **1**-**2** may be linked to specific targets or receptors within AT. Once triggered, these compounds likely initiate downstream signaling pathways that are tailored to the corresponding interactions, ultimately leading to the observed phenotypic effects during our experiments. Future work will seek to investigate more details for these mechanisms.

One of the objectives of this study is to elucidate the impact of secondary metabolites produced by G7 and, based on this, to explain their inhibitory effect on plant pathogens and beneficial role for crops. Notably, the treatment with G7 resulted in significant phenotypic changes, such as shorter primary roots and increased root hair density. A plausible explanation for this phenomenon is a decrease in the cell division capacity of root tip cells, leading to premature differentiation of cells in the mature zone, resulting in the formation of root hairs. The growth of primary roots is tightly regulated by various factors, including hormone biosynthesis, signalling, and transport, with auxin playing a particularly important role. The precise distribution patterns and polar transport of auxin are controlled by various influx and efflux carriers, such as AUX/LAX and PIN proteins, as well as intracellular carriers like PILs. These processes ensure the regulatory function of auxin^[21]^. In this study, we employed a multi-omics approach to rapidly identify the affected pathway and establish that the auxin concentration gradient was significantly impacted under G7 treatment since QC homeostasis. A significant reduction in the abundance of PIN2, PIN3, PIN4 and PIL3 proteins was discerned within the proteomic analysis, thereby implying that the perturbation of the auxin concentration gradient could be attributed to a decrement in the polar transport efficacy of auxin. As depicted in figure 6, the influence of lydicamycin **1** treatment on the distribution of auxin, and the localization and expression of the PIN protein family becomes apparent. Notably, treatment with **1** induces a substantial repositioning of PIN2 and effectively suppresses the fluorescence signal emitted by *PIN1::PIN1-GFP, PIN4::PIN4-GFP*, and *PIN7::PIN7-GFP*.

Finally, we acknowledge that there are still certain complexities that require further investigation in future research. One such complexity is the complete identification of the specific targets or receptors of lydicamycins. Despite this, our work presents a comprehensive model of plant-microbe interaction, illustrating a complex molecular regulatory network pattern as shown in figure 7. This model has the potential to enhance our understanding of the impacts of microbial natural products on plant growth and development across various scales and dimensions, including the herbicides discussed in this study. Moreover, as evidenced in our research, the re-evaluation of previously overlooked natural products has revealed intriguing and unexpected biological activities. The isolation of well-known natural product families, particularly those exhibiting diverse congeners within a stable ecosystem, can hold significant practical implications, as demonstrated by our findings. Therefore, exploiting unexplored plant-microbe ecosystems paves a useful way for natural product discovery, regardless of whether they are novel or known.

**Figure 7.**
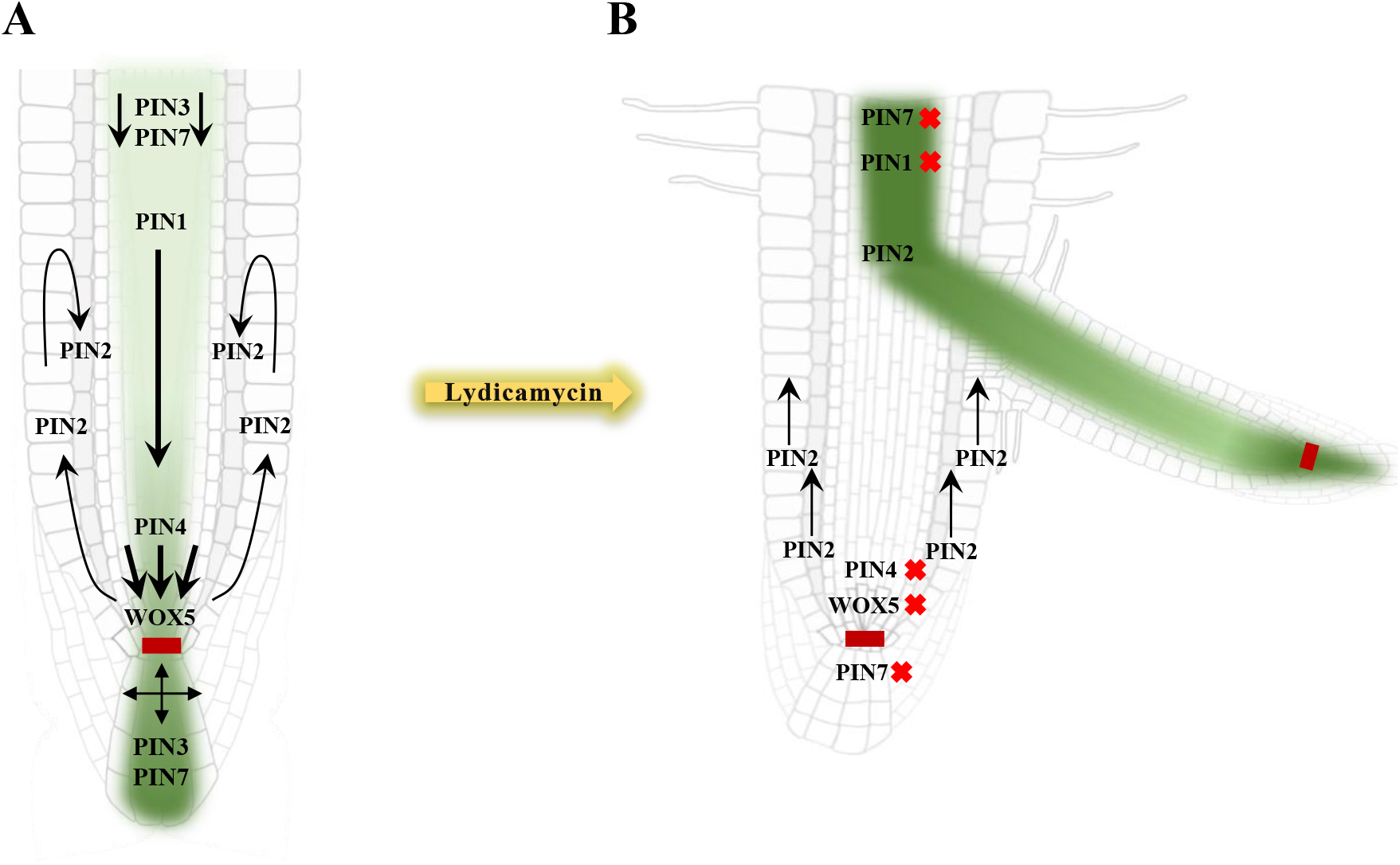
A proposed model in this study presents an overview of the spatial distribution of PIN proteins and auxin within the AT root tip under lydicamycin treatment. Schematic diagram A illustrates the specific localization patterns of PIN proteins and the distribution of auxin within the root meristem. The green color gradient indicates varying levels of auxin concentration in the root tip. Notably, PIN1 and PIN4 proteins play a crucial role in facilitating acropetal auxin flow in the root stele, as depicted by the black arrows. In contrast, PIN2 mediates basipetal auxin transport through the root epidermis, as indicated by the black arrows. PIN3 and PIN7 show prominent expression in root columella cells. The red rectangle represents the Quiescent Center. Schematic diagram B illustrates the aberrant localization patterns of PIN proteins and the disrupted distribution of auxin within the root meristem under lydicamycin treatment. Remarkably, the expression levels of PIN1, PIN4, and PIN7 appear to be either unexpressed or significantly reduced compared to their regular distribution, resulting in the QC disorder. Furthermore, the distribution of PIN2 shows alterations, indicating a deviation from the expected basipetal auxin flow through the root cortical cells. This disruption of the auxin gradient distribution exerts a pronounced impact on the subsequent lateral root generation.

## Materials and methods

### Standard materials and methods

*Arabidopsis thaliana* ecotype Col-0, along with several transgenic marker lines including *DR5::GFP, CYCB1;1pro:GUS, WOX5::GFP, PIN1::PIN1-GFP, PIN2::PIN2-GFP, PIN4::PIN4-GFP* and *PIN7::PIN7-GFP* were employed as experimental materials in this study and were graciously provided by Prof. Haiyun Ren and Ting Wang from Beijing Normal University. The rice wild-type *japonica* (*O*.*sativa*) cultivar was was kindly supplied by Prof. Xiaoyan Wang.

Unless otherwise stated, all chemicals were provided by Shanghai Macklin Biochemical Co., Ltd. All organic solvents used were of HPLC grade or of equivalent quality. Samples were analyzed by LC-MS/MS using an Agilent G6500 UPLC system coupled to a quadrupole time-of-flight mass spectrometer. The instrument was calibrated using the HP-0921 ion (calculated mass-to-charge ratio 922.0098) following the manufacturer’s instructions and operated in positive-negative mode switching. The following LC-MS/MS and UPLC methods were employed throughout the study unless indicated otherwise: a Phenomenex Kinetex C_18_ column (100 × 2.1 mm, 100 Å), mobile phase A: water with 0.1% formic acid, and mobile phase B: acetonitrile with 0.1% formic acid. The elution gradient was as follows: 0-1 min, 20% B; 1-12 min, 20-100% B; 12-14 min, 100% B; 14-14.1 min, 100-20% B; 14.1-17 min, 20% B. The flow rate was set at 0.3 mL min^−1^, and the injection volume was 10 μL. For solid agar medium samples, a rectangular piece of agar (1 cm^3^) was extracted by shaking it in methanol (1 mL) for 5 minutes. The resulting crude extract was transferred to a clean tube, and the solvent was evaporated under reduced pressure. The obtained extract was reconstituted in methanol (100 μL) for further analysis.

### Plant growth conditions

To initiate the experimental procedure, seeds were subjected to surface sterilization using 50% ethanol solution for 5 min, followed by treatment with 0.5% NaOCl for an additional 5 min. Subsequently, the sterilized seeds were rinsed three times with water. Afterward, the seeds were sown on ½ MS medium, and then placed in darkness at a temperature of 4°C for 2 d. The seeded petri dishes were then vertically positioned in a growth chamber, set to a long-day photoperiod of 16 hours of light with a light intensity of 300 μM m^−2^ s^−1^, while maintaining a constant temperature of 22°C as preset growth conditions.

### Plant-microbe cocultivation

The spores suspension (OD_600_ = 0.6) of G7 and G7_*Δlyd67* was spread at a 3×10 cm (for AT seedlings) or 3×13 cm (for rice seedlings) on the ½ MS gel medium area at a square petri dish (10×10 cm or 13×13 cm). After 4 d of fermentation, a set of 2 -day-old seedlings were transferred and grew in preset conditions. Specifically, the AT seedlings were grown for 4 d, whereas 13 d for rice.

### Plant phenotypic analysis

To measure shoot and root fresh weight, 6 d AT or 13 d rice seedlings were cut at the root-shoot junction. The separated shoots and roots were divided into three groups, with each group containing 15 or 10 samples, respectively. The weight of each group was immediately measured to determine their fresh weights accurately. For assessing root length, lateral root number (greater than 0.5 mm), and total root number, the digital images of seedlings in Petri dishes were captured using a Canon ES-71II digital camera. The AT primary root length was measured precisely using Digimizer version 4.5. To determine the size of the root meristematic zone and the number of cells, confocal images of PI-stained roots from seedlings were analyzed using Digimizer version 4.5. The root meristematic zone size was calculated by counting the cortical cells between the quiescent center and the first cell that was twice the length of the preceding cell. Root hairs were observed with ZEISS Axio Zoom.V16.

### Confocal imaging analysis

High-resolution microscopic images were acquired using the ZEISS confocal microscope system Lsm980. The roots of 6-day-old seedlings were stained with propidium iodide (PI) at a concentration of 5 µg·mL −1 for 10 s. The excitation wavelength used was 587 nm, and the detection window was set at 550-700 nm. For the 6-day-old seedlings expressing *DR5::GFP, WOX5::GFP, PIN1::PIN1-GFP, PIN2::PIN2-GFP, PIN4::PIN4-GFP*, and *PIN7::PIN7-GFP* marker lines, the chromophores were excited at a wavelength of 488 nm, and fluorescence emission was detected within the range of 492-562 nm.

### GUS histochemical staining

Histochemical staining for GUS activity was conducted on 6-day-old *CYCB1;1pro:GUS* seedlings. The staining procedure involved the use of a staining solution composed of 50 mM sodium phosphate (pH = 7), 10 mM EDTA, 0.5 mM K_4_[Fe(CN)_6_], 0.5 mM K_3_[Fe(CN)_6_], 0.5 mM 5-bromo-4-chloro-3-indolyl-β-glucuronic acid, and 0.01% Silwet L-77. The seedlings were submerged in the staining solution and incubated at 37°C overnight. Following GUS staining, the roots were treated with a clearing solution consisting of chloral hydrate, glycerol, and water in an 8:1:2 ratio. The cleared roots were then examined using a ZEISS Axio Imager Z2 microscope.

### Production and isolation of lydicamycins

The pure compounds of lydicamycins used in this study were isolated previously, for detailed protocols regarding the microbial fermentation and compound purification, please see our recent work^[8]^.

### Herbicidal activity test of lydicamycins

After 2 d of growth, the seedlings were carefully transferred to 96-well plates. Each well of the plate was filled with 120 µL of ½ MS liquid medium containing lydicamycin compounds with concentration range 10, 20, 40, 60, 80, 100 and 200 μM, respectively. The seedlings were then grown in the preset conditions for 4 d before taken into investigation.

### Bioactivity test

The aforementioned strains were cultured on ISP2 agar at a temperature of 28 °C for a duration of 7 days. To assess their antimicrobial activity, commonly used laboratory strains such as *Bacillus subtilis, Escherichia coli*, and *Candida albicans* were employed as indicators. Additionally, several phytopathogenic fungi including *Botrytis cinerea, Fusarium graminearum, Ustilaginoidea virens*, and *Magnaporthe grisea* were used for antifungal screening. The screening was conducted using the agar plug assay. Initially, 200 µL of fresh liquid cultures of *B. subtilis, E. coli*, and *C. albicans* were streaked onto solid LB agar plates. Then, agar plugs containing bioactive compounds from the incubated ISP2 medium were collected using sterile 1 mL pipette tips with round ends. These agar plugs were placed on LB solid medium and allowed to incubate with the test strains. The presence of bioactive compounds produced by the rhizobacteria was confirmed by the observation of apparent inhibition zones after 12 hours. The antifungal activity test was performed in a similar manner except that PDA solid medium was used for fungal incubation. Each bioassay test was repeated independently for a minimum of three times to ensure reliability and consistency of results.

### Multiomics analysis under G7 treatment

Multi-omics analysis was conducted on samples from the TG7 treatment group and the mock-treated group. All testing samples were collected from three independent experiments, with multiple plates used within each experiment to ensure robustness and reproducibility.

For transcriptomic analysis, the cleaned RNA-seq reads were aligned to the *Arabidopsis thaliana* genome (TAIR50) using the HISAT2 v2.2.1^[22]^. The mapped reads were then quantified using featureCounts^[23]^. Differential expression analysis was performed on the quantified reads and protein counts using the DESeq2 R package. Genes with a count-per-million (CPM) value greater than 0.5 in at least two samples were included in the analysis of differentially expressed genes (DEGs). DEGs in the TG7 group were identified based on a fold change of |log2FC| > 0.58 and a false discovery rate (FDR) of less than 0.05, using a Student’s t-test with Benjamini-Hochberg correction.

For proteomics analysis, differentially expressed proteins (DEPs) were identified based on the criteria of |log2FC| > 0.26 and FDR < 0.05. To gain further insights into the functional implications of the identified DEGs and DEPs in the TG7 group, gene ontology (GO) biological processes enrichment analyses were performed. The downregulated and upregulated DEGs and DEPs were analyzed simultaneously using the online GO enrichment server PANTHER (http://geneontology.org/)^[24]^. To reduce redundancy, the results were further processed using REVIGO (http://revigo.irb.hr/)^[25]^.

For metabolomic analysis, all multivariate data analyses and modelling were performed using Ropls software^[26]^. Principal component analysis (PCA), orthogonal partial least squares discriminant analysis (PLS-DA), and partial least squares discriminant analysis (OPLS-DA) were employed to build the models. To assess the robustness of the models, permutation tests were performed to evaluate potential overfitting. The OPLS-DA model allowed the identification of discriminating metabolites using the variable importance on projection (VIP). The significance of metabolite contributions for classification was determined using the *p* values, VIP values, and fold change (FC). Differentially expressed metabolites (DEM) with *p* < 0.05 and VIP values > 1 were considered statistically significant.

### RNA isolation and quantitative real-time PCR analysis

The total RNA for each sample was extracted with the plant total RNA extraction kit (ELK Biotechnology). Complementary DNA was synthesized from RNA by using HiScript® III 1st Strand cDNA Synthesis Kit (+gDNA wiper) (Vazyme). Real-time PCR was performed using ChamQ Universal SYBR qPCR Master Mix (Vazyme) on the QuantStudio 6 Flex (Life Technologies). Briefly, the qPCR program run consisted of a first step at 95°C for 30 s and afterwards 40 cycles alternating between 10 s at 95°C and 30 s at 60 °C ^[27]^. The internal reference, *ACTIN2*, was adopted to normalize the expression levels of the target genes. The specific gene primers were designed by Primer Premier 5.0 and were listed in Supplementary Table 1.

### Statistical analysis

Quantitative data were expressed as means ± SD. The statistical significance of the experimental data was assessed using Student’s t-test, and significant differences were denoted by asterisks (* *p* ≤ 0.05, ** *p* ≤ 0.01, and *** *p* ≤ 0.001).

## Author Contributions

Z.Q. and H.Z. designed and supervised the research. J.H. performed detailed bioinformatic analysis and the physiological experiments, X.L. performed the chemical experiment, J.L. analysed the mechanisms. All authors analysed and discussed the data. H.Z., J.H. and Z.Q. wrote the manuscript and all authors commented.

## Acknowledgements

This work was supported by the National Natural Science Foundation of China (32170079 to Z.Q. and 32200035 to H.Z.), the Natural Science Foundation of Guangdong (2021A1515012026 to Z.Q.), Guangdong Talent Scheme (2021QN020100 to Z.Q.), Guangdong Innovation Research Team for Plant-Microbe Interaction (2021KCXTD037 to Z.Q.), as well as Beijing Normal University via the Youth Talent Strategic Program Project (310432104 to Z.Q.). S.P. and X.L. thank the Postgraduate Interdisciplinary Project from Beijing Normal University.

## Conflicts of interest

The authors declare no competing financial interests.

## Notes

### Competing Interest Statement

The authors have declared no competing interest.

