## Supplemental information for "Extracting herbicide and antibiotic natural products from a plant-microbe interaction system"

**Correspondence**

**Supplementary Figures**

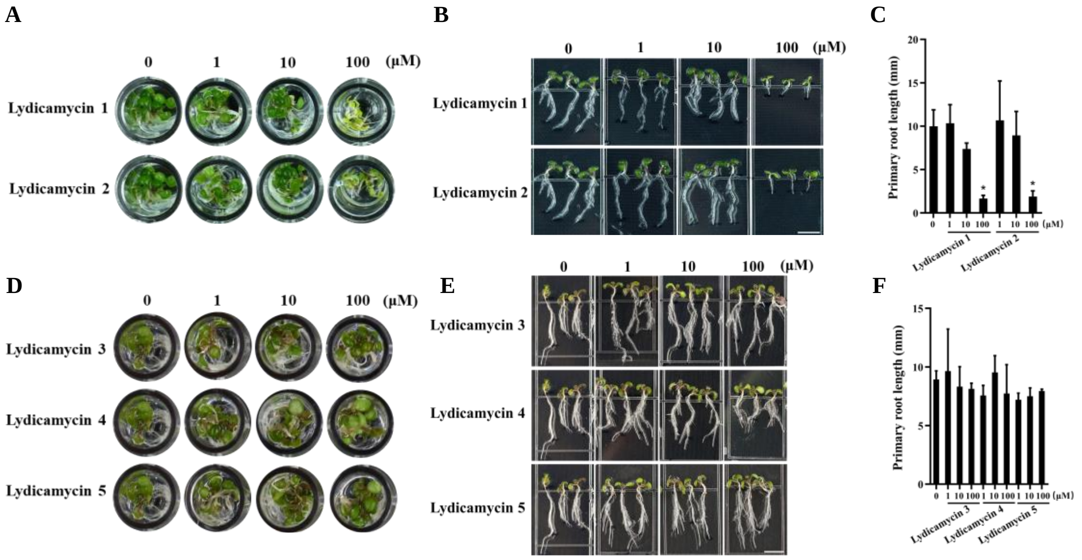


**Figure S1.** The effects of lydicamycins on primary root growth of AT seedlings. AT Seedlings were grown on ½ MS liquid medium containing various concentrations of **1**-**2** (A) and **3-5** (D), with water as the solvent. (B and E) Representative images of AT seedlings from panel A and D. Scale bar: 5 mm. (C and F) Quantification of primary root length in panel A and D. The data represents the mean ± SD of 20-50 seedlings (t-test, *, p<0.05). The experiment was repeated twice with similar results.

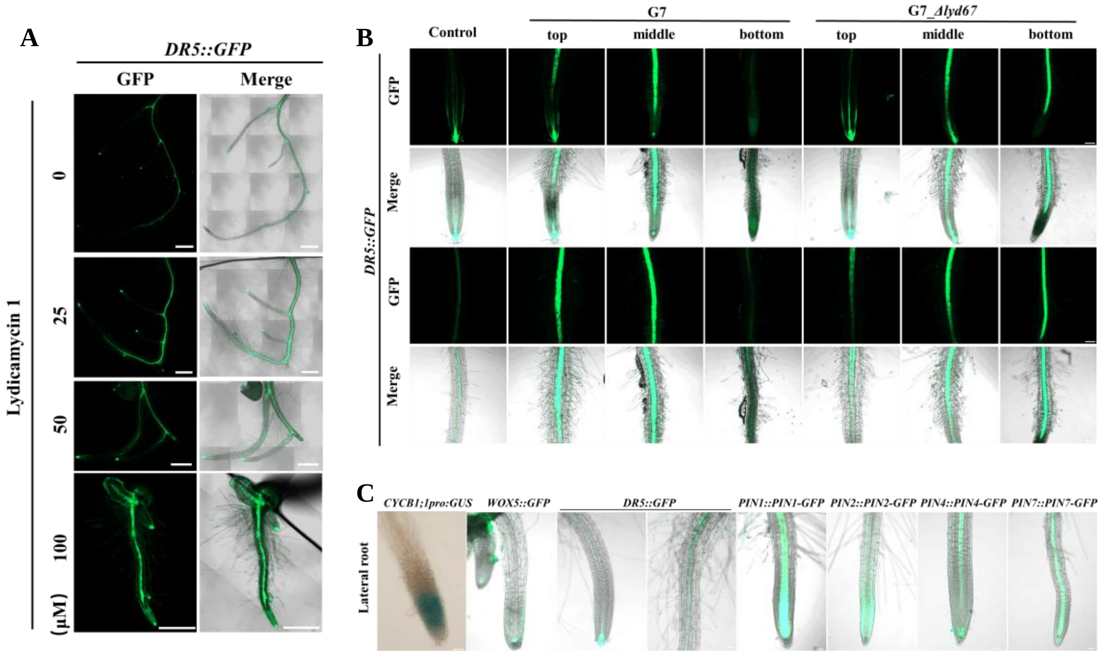


**Figure S2.** The effects of lydicamycins produced by G7 on auxin transport in AT primary root tips. (A) The representative images of *DR5::GFP* AT cultivated in the medium supplemented with various concentrations of **1**. Scale bar: 500 µm. (B) The representative images of *DR5::GFP* AT seedlings cultivated with G7 or G7_*Δlyd67*. Scale bar: 100 µm. (C) Representative images of *CYCB1;1pro:GUS*, *WOX5::GFP*, *DR5::GFP*, *PIN1::PIN1-GFP*, *PIN2::PIN2-GFP*, *PIN4::PIN4-GFP*, and *PIN7::PIN7-GFP* AT seedlings treated with 50 µM **1**. The images show lateral roots abnormally protruding from the primary root. Scale bar: 100 µm.

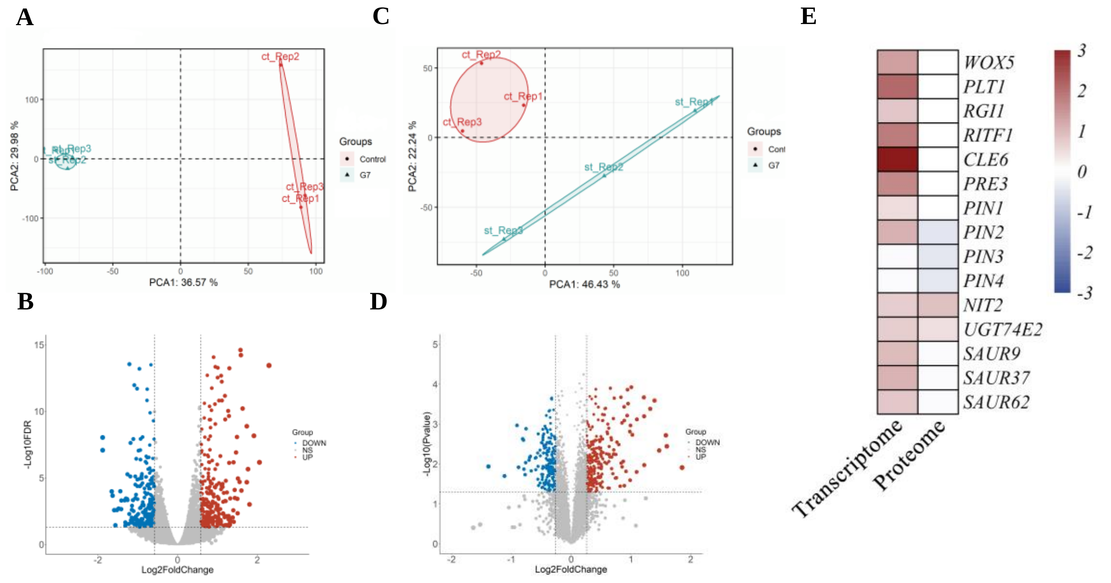


**Figure S3.** The multiomics analysis. (A) PCA of transcriptomes with or without G7 treatment represented in a two-dimensional space. (B) Volcano plot showed the differentially expressed genes between G7 and untreated AT. (C) PCA of proteomes with or without G7 treatment represented in a two-dimensional space. (D) Volcano plot showed the differentially expressed proteins between G7 and untreated AT. (E) Heatmap plot showed QC homeostasis and IAA metabolism, transport, and signal transduction differentially expressed associated genes and proteins in transcriptomes and proteome.

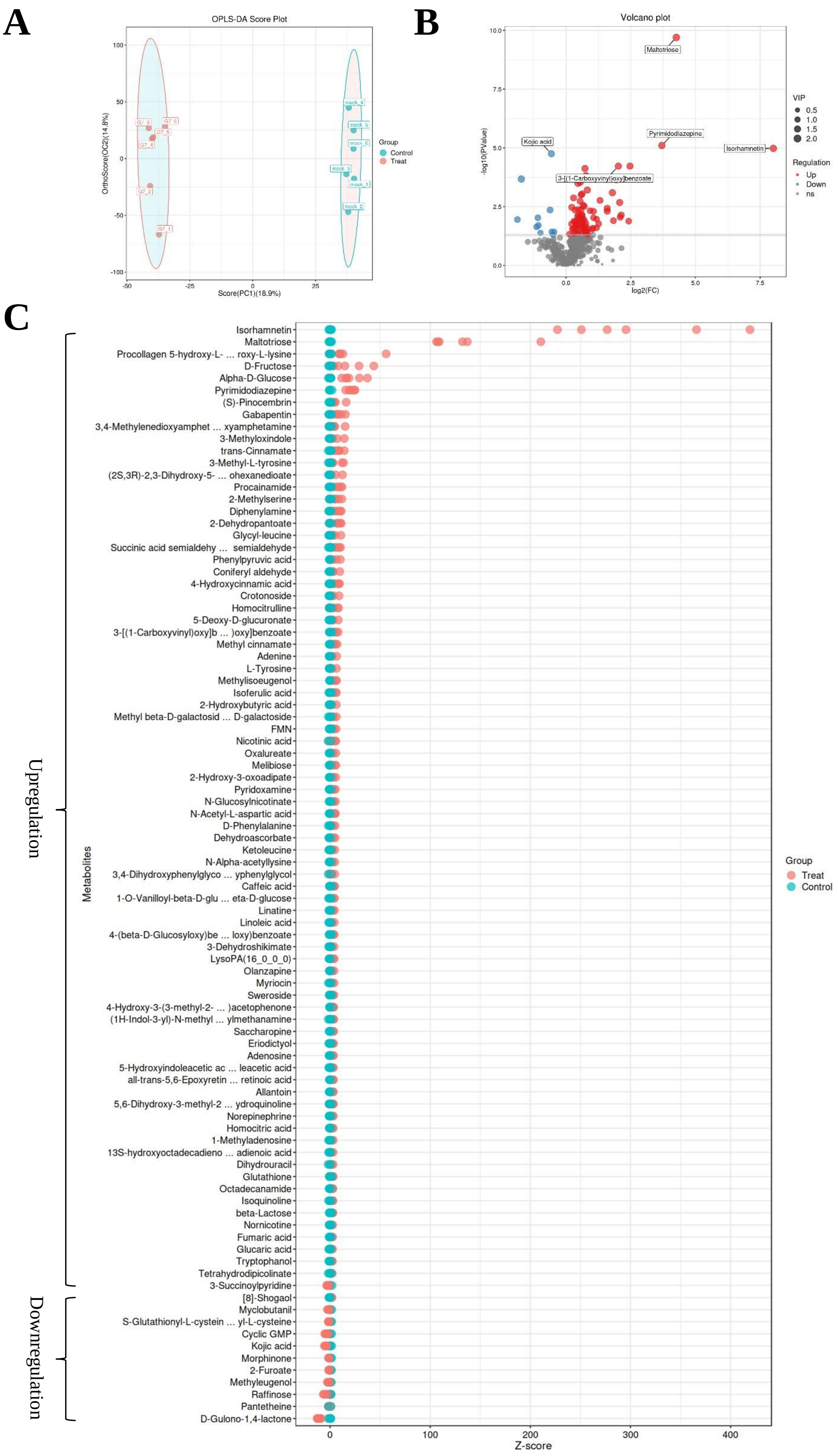
A
OPLS-DA Score Plot
OrthoScore(OC2)(14.8%)
Score(PC1)(18.9%)
Group
Control
Treat
B
Volcano plot
-log10(PValue)
log2(FC)
Maltotriose
Pyrimidodiazepine
Isorhamnetin
Kojic acid
3-[(1-Carboxyvinyl)oxy]benzoate
VIP
0.5
1.0
1.5
2.0
Regulation
Up
Down
ns
C
Upregulation
Downregulation
Metabolites
Isorhamnetin
Maltotriose
Procollagen 5-hydroxy-L- ... roxy-L-lysine
D-Fructose
Alpha-D-Glucose
Pyrimidodiazepine
(S)-Pinocembrin
Gabapentin
3,4-Methylenedioxyamphet ... xyamphetamine
3-Methyloxindole
trans-Cinnamate
3-Methyl-L-tyrosine
(2S,3R)-2,3-Dihydroxy-5- ... ohexanedioate
Procainamide
2-Methylserine
Diphenylamine
2-Dehydropantoate
Glycyl-leucine
Succinic acid semialdehy ... semialdehyde
Phenylpyruvic acid
Coniferyl aldehyde
4-Hydroxycinnamic acid
Crotonoside
Homocitrulline
5-Deoxy-D-glucuronate
3-[(1-Carboxyvinyl)oxy]b ... )oxy]benzoate
Methyl cinnamate
Adenine
L-Tyrosine
Methylisoeugenol
Isoferulic acid
2-Hydroxybutyric acid
Methyl beta-D-galactosid ... D-galactoside
FMN
Nicotinic acid
Oxalureate
Melibiose
2-Hydroxy-3-oxoadipate
Pyridoxamine
N-Glucosylnicotinate
N-Acetyl-L-aspartic acid
D-Phenylalanine
Dehydroascorbate
Ketoleucine
N-Alpha-acetyllysine
3,4-Dihydroxyphenylglyco ... yphenylglycol
Caffeic acid
1-O-Vanilloyl-beta-D-glu ... eta-D-glucose
Linatine
Linoleic acid
4-(beta-D-Glucosyloxy)be ... loxy)benzoate
3-Dehydroshikimate
LysoPA(16_0_0_0)
Olanzapine
Myriocin
Sweroside
4-Hydroxy-3-(3-methyl-2- ... )acetophenone
(1H-Indol-3-yl)-N-methyl ... ylmethanamine
Saccharopine
Eriodictyol
Adenosine
5-Hydroxyindoleacetic ac ... leacetic acid
all-trans-5,6-Epoxyretin ... retinoic acid
Allantoin
5,6-Dihydroxy-3-methyl-2 ... ydroquinoline
Norepinephrine
Homocitric acid
1-Methyladenosine
13S-hydroxyoctadecadieno ... adienoic acid
Dihydrouracil
Glutathione
Octadecanamide
Isoquinoline
beta-Lactose
Nornicotine
Fumaric acid
Glucaric acid
Tryptophanol
Tetrahydrodipicolinate
3-Succinoylpyridine
[8]-Shogaol
Myclobutanil
S-Glutathionyl-L-cystein ... yl-L-cysteine
Cyclic GMP
Kojic acid
Morphinone
2-Furoate
Methyleugenol
Raffinose
Pantetheine
D-Gulono-1,4-lactone
Z-score
Group
Treat
Control

**Figure S4.** The metabolome analysis. (A) Orthogonal Projections to Latent Structures Discriminant Analysis (OPLS-DA) of metabolome with or without G7 treatment represented in a two-dimensional space. (B) Volcano plot showed the differentially expressed annotated metabolites with or without G7 treatment. (C) Bubble plot showed the Z-score of differentially expressed annotated metabolites with or without G7 treatment. The horizontal axis depicts the numerical values derived from the conversion of the relative content of metabolites within the group into Z-scores. As one moves towards the right on the axis, it signifies an increasing abundance of the respective metabolite within that particular group.

**Supplementary Tables**

Note that due to the scales Table S2-3 are deposited as independent Excel spreadsheets.

**Table S1.** Primers used in this study.

| Gene ID | Gene name | Primer Sequence (5' → 3') |
|---|---|---|
| AT3G11260 | *WOX5* | F: GCAACAATAACGGAGGAACG<br>R: AGACCGGCTCGAAACAGATC |
| AT3G20840 | *PLT1* | F: AGTAGCACTCTTCCCATCGG<br>R: ACCAAGGGCTATCATCTCCG |
| AT1G51190 | *PLT2* | F: TGGCTGATTTCTTAGGAGTG<br>R: TTAGGAGCATTAGAGGCACA |
| AT2G04025 | *RGF3* | F: TGTTTCTCCTTTCGCTATTC<br>R: TCAGTTTTCGTCCTCGTATC |
| AT3G30350 | *RGF4* | F: AAAAGAAGAGAGCATAAGCAGACC<br>R: AATCATCACGCAACAACCGA |
| AT3G24240 | *RGI1* | F: TGGGTGAGGCAGAATAGAGG<br>R: CACACACAGCAAAGCAGTGC |
| AT5G48940 | *RGI2* | F: CCAGAATACGGATACTCAAT<br>R: CTTGCTTGTAGTCCTTGGTC |
| AT2G12646 | *RITF1* | F: ATCCATTGCTCCTTGGGTTG<br>R: CCGCCACTTCATCATCTGTC |
| AT2G31085 | *CLE6* | F: GCTCGAATCCTCCGTACATA<br>R: AAAAACCGTCTCTCGTCTTG |
| AT1G74500 | *PRE3* | F: CGAGGCAATCATCAGGAACT<br>R: TGAAACCTTGTCGGAACGAC |
| AT1G73590 | *PIN1* | F: CAAATCGTTGTTCTTCAGTG<br>R: TTCCAAAGGTTGTCTTCCAT |
| AT5G57090 | *PIN2* | F: CGTTATCCTCGCCGCACTCT<br>R: TCCGTACATCGCCCTAAGCA |
| AT1G70940 | *PIN3* | F: ATACGAATCAGCAGACGACT<br>R: CGGATCTCTTTAGCACCTTG |

| | | |
|---|---|---|
| AT2G01420 | *PIN4* | F: GGCTAACCTAACCAAGAACG<br>R: TGGACCATTAGAGAACCTGC |
| AT4G25960 | *ABCB2* | F: AACCCTTCCTTGAATCGCAC<br>R: TTTTTGGATGGATCAGCCCC |
| AT1G77690 | *LAX3* | F: CGGCAGAGAAAATAGAGACA<br>R: ACTAAACCAAGCATCGTAGA |
| AT1G76520 | *PILS3* | F: CTGCTTACATTCATCATTGG<br>R: GGGACCTCCTTTTTCTTTAC |
| AT1G04610 | *YUCCA3* | F: CCTTGAGTCCTACGCAGCCA<br>R: CCGTTGCCACCACAATCCAT |
| AT1G70560 | *TAA1* | F: GCCGCCGCTCCTTTTTACTC<br>R: TGATGGTTCCGTCAGGGTTA |
| AT3G44300 | *NIT2* | F: ACAACGATACTCCCGCCACT<br>R: GAACTCCCACCCCTAAACCA |
| AT1G05680 | *UGT74E2* | F: CTCACTCTGGTCCTCGTCTC<br>R: TCTTGTAATGGTTCCTCGCC |
| AT2G23170 | *GH3.3* | F: AAGGGATTCACTGACCGTAA<br>R: GATGGGGTAAGAAGACAAGA |
| AT1G29430 | *SAUR62* | F: ATCTGAAGAAGAGTTCGGTC<br>R: GCACTACAATGTTGTTGTGG |
| AT4G31320 | *SAUR37* | F: TGTTGCGTGAAGCAGAAGAA<br>R: TCCGAAGGATGACGATAACT |
| AT3G23050 | *IAA7* | F: TTGAGAGTCCTGCCAAATCG<br>R: GTCTTCTCCTTGGGAACAGC |
| AT3G15540 | *IAA19* | F: GGACTCGGGCTTGAGATAAC<br>R: TCACCACCAGATGAAACGAC |
| AT3G18780 | *ACTIN2* | F: GGTAACATTGTGCTCAGTGGTGG<br>R: AACGACCTTAATCTTCATGCTGC |
